# CasRx-mediated RNA targeting prevents choroidal neovascularization in a mouse model of age-related macular degeneration

**DOI:** 10.1101/2020.02.18.955286

**Authors:** Changyang Zhou, Xinde Hu, Cheng Tang, Wenjia Liu, Shaoran Wang, Yingsi Zhou, Qimeng Zhao, Qiyu Bo, Linyu Shi, Xiaodong Sun, Haibo Zhou, Hui Yang

**Author notes:** These authors contributed equally to this work (Changyang Zhou, Xinde Hu, Cheng Tang, Wenjia Liu, Shaoran Wang).

## Abstract

The smallest Cas13 family protein, CasRx, has a high cleavage activity and targeting specificity, offering attractive opportunity for therapeutic applications. Here we report that delivery of CasRx by adeno-associated virus via intravitreal injection could efficiently knockdown *Vegfa* transcripts and significantly reduce the area of laser-induced choroidal neovascularization in a mouse model of age-related macular degeneration. Thus, RNA-targeting CRISPR system could be used for *in vivo* gene therapy.

## Main

Age-related macular degeneration (AMD), characterized by the development of choroidal neovascularization (CNV), is a leading cause of vision deterioration in adults over age 50^1^. An angiogenic growth factor vascular endothelial growth factor A (VEGFA) plays a crucial role in CNV pathogenesis and anti-VEGFA therapy using humanized antibodies has been widely used in treating AMD, with the therapeutic effects maintained by regular injections of antibodies^2,3^. Two recent studies showed that in a mouse model of AMD that permanent *Vegfa* gene disruption could be induced by SpCas9 or LbCpf1 editing^4,5^. However, risks associated with permanent DNA modifications, including unwanted off-target and on-target effects, need to be considered^6,7^.

The Cas13 protein family was recently shown to be a programmable RNA-targeting CRISPR system^8–13^, which could mediated RNA knockdown with high efficiency and specificity relative to other existing RNA interference approaches^8,9,12^. Several Cas13 proteins have been identified, among which RfxCas13d (CasRx) has the smallest size and highest RNase activity^12^. Here, we examine the potential application of CasRx system for *in vivo* gene therapy, using a laser-induced mouse model of AMD. Our results show that adeno-associated viral (AAV)-delivered CasRx could knockdown *Vegfa* transcripts efficiently, resulting in the significant reduction of CNV area in this AMD model.

We first identified two CasRx targeting sites that are conserved in the human and mouse *Vegfa* gene. To achieve efficient *Vegfa* mRNA knockdown, two guide RNAs (gRNAs) targeting these two sites respectively were designed (Fig. 1a). We found that transient transfection of vectors expressing CasRx and the gRNA resulted in marked reduction of the *Vegfa* mRNA level in cultured human 293T cells (12+/− 3.5%, s.e.m.) and mouse N2a cells (29.5 +/− 8.4%, s.e.m.) within two days, as compared to cells transfected with the control vector (Fig. 1b, c). The VEGFA protein levels were also significantly reduced in mouse N2a cells (Fig. 1d). To determine targeting specificity of CasRx, we performed transcriptome-wide RNA-seq analysis. Besides Vegfa, the expression levels of many other genes were changed and more than half of top-ranked genes with altered expression were related with *Vegfa* according to previous studies. (Supplementary Fig. 1 and Supplementary Table 1). To investigate the knockdown efficiency of CasRx in the normal mouse retina, we intravitreally injected AAVs encoding CasRx and a dual-gRNA array targeting *Vegfa* (referred as AAV-CasRx-*Vegfa*). Three weeks after injection, choroid-retinal pigment epithelial (RPE) tissue complex was isolated for qPCR analysis (Fig. 1e,f). We observed the expression of AAV-CasRx-*Vegfa* (Fig. 1g) and found that *Vegfa* transcripts in the treated eye were potently suppressed (65.4 +/− 8.7%, s.e.m), as compared to those in the contralateral eye injected with PBS (Fig. 1h).

**Figure 1.**
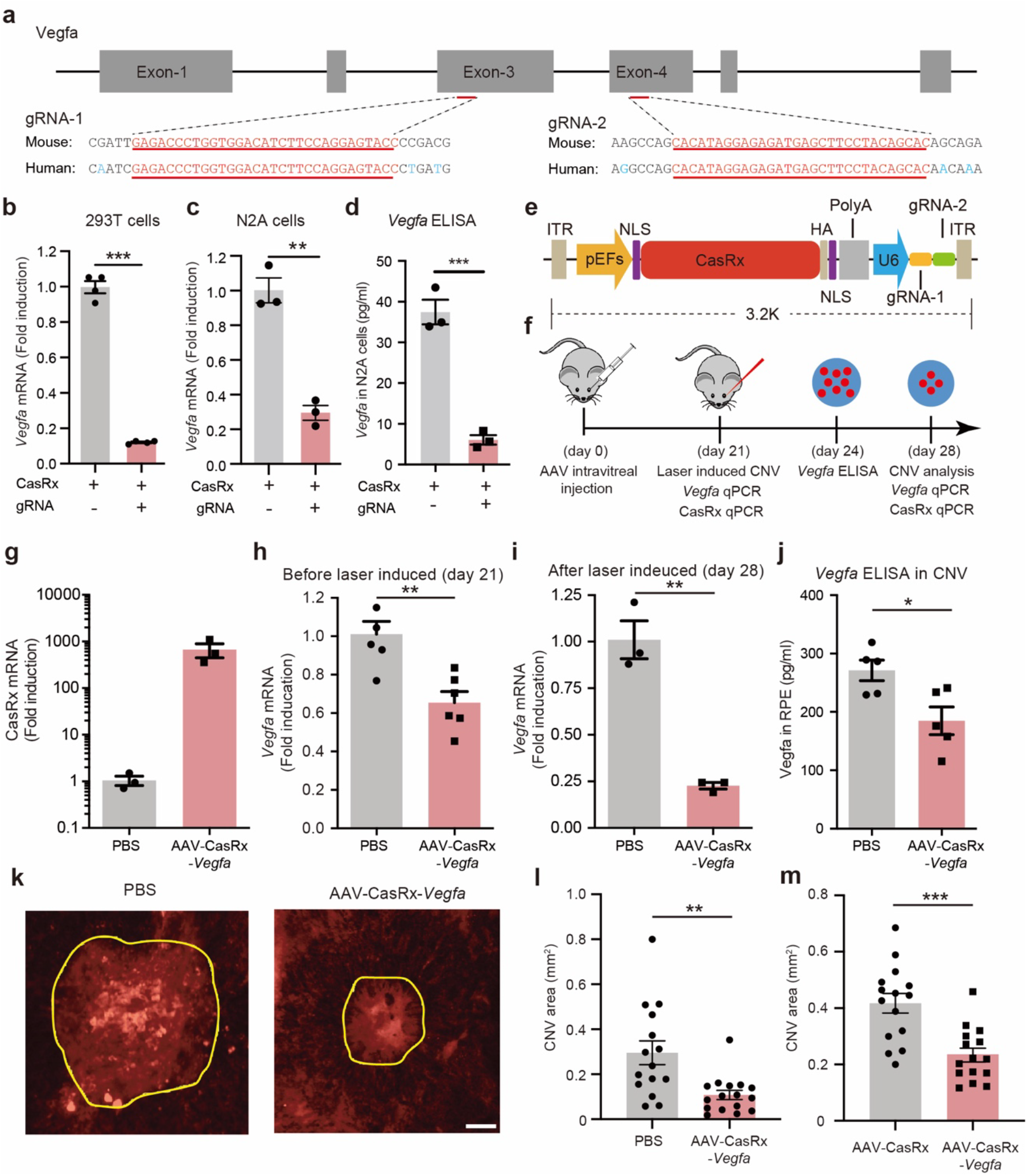
AAV-mediated delivery of CasRx reduces the area of CNV in a mouse model of AMD. (**a**) Schematic illustration of the targeting sites. The CasRx targeting sites are conserved in the human and mouse *Vegfa* gene, and all isoforms were targeted. (**b, c)** Transient transfection of AAV vectors can potently knock down *Vegfa* in both human 293T (*n* = 4 repeats, *p* < 0.0001, *t* = 25.02) cells and mouse N2a cells (*n* = 3 repeats, *p* = 0.0011, *t* = 8.425). (**d**) VEGFA protein levels (*n* = 3 repeats, *p* < 0.01, t =9.675). (**e**) Schematic showing AAV-CasRx-*Vegfa.* (**f**) Schematic of the experimental procedure. AAV-CasRx-*Vegfa* was intravitreally injected into one eye, and AAV-CasRx was injected into the other eye as a control, 21 days before laser burn. Three weeks after AAV infection, the transcription level of *Vegfa* mRNA was analyzed without laser burn. VEGFA protein levels were quantified by ELISA 3 days after laser burn. CasRx and *Vegfa* mRNA levels as well as the area of CNV were measured 7 days after laser burn. (**g**) CasRx mRNA levels without laser burn, 21 days after AAV injection (*n* = 3 mice). (**h,i)** *Vegfa* mRNA levels before or 7 days after laser burn (Before laser burn: *n* = 6 mice, *p* = 0.002, *t* = 4.059; after laser burn: *n* = 3 mice, *p* = 0.002, *t* = 7.583). (**j)** VEGFA protein levels 3 days after CNV induction (*n* = 5 mice, *p* = 0.019, *t* = 2.928). (**k**) Representative CNV images injected with the PBS or AAV-CasRx-*Vegfa*, 7 days after laser burn. The area of CNV is indicated by the yellow line. Scale bar: 200 μm. (**l,m**) The CNV area. A data point represents a laser burn and in total 4 laser burns were induced in each eye. (PBS + AAV-CasRx-*Vegfa*: *n* = 4 mice, *p* = 0.002, *t* = 3.39; AAV-CasRx + AAV-CasRx-*Vegfa*: *n* = 4 mice, *p* = 0.0002, *t* = 4.292). All values are presented as mean ± s.e.m.. \**p* < 0.05, \*\**p* < 0.01, \*\*\**p* < 0.001, unpaired t-test.

We next created the AMD mice by inducing CNV in both eyes by laser irradiation (Supplementary Fig. 2a,b, also see Methods). To investigate the potential usefulness of mRNA knockdown approach for treating AMD, we injected AAV-CasRx-*Vegfa* into one eye of the mouse, and PBS in the other eye as control (Fig 1f. Induction of CNV was performed in both eyes three weeks later. After laser burn, we confirmed successful infection of AAV-CasRx-*Vegfa* (Supplementary Fig. 3a). Furthermore, we found that the levels of *Vegfa* mRNA and VEGFA protein were significantly lower in the AAV-injected eye as compared that those in the contralateral PBS-injected eye (mRNA, 22.7 +/− 1.8% s.e.m., *p* = 0.002; protein, 68.2 +/− 8.7%, s.e.m., *p* = 0.019; unpaired *t*-test) (Fig. 1i,j). Thus, intravitreal injection of the *Vegfa* mRNA-targeting AAV was efficient to knockdown VEGFA expression in the injected eye. The therapeutic effect of this CasRX approach was assessed by quantifying the CNV area 7 days after laser treatment. Our results showed that *Vegfa*-targeting AAV markedly reduced the area of CNV at two different levels of laser irradiation, as compared to the control eyes injected with PBS (Fig. 1k,l and Supplementary Fig. 3b and 4a,b. 180 mW, 66 +/− 7.8%, s.e.m., *n* = 6 mice, *p* = 0.004; 240 mW, 36.5 +/− 6.9%, s.e.m., *n* = 4 mice, *p* = 0.002; unpaired *t*-test). Reduction of CNV was also confirmed by injecting AAV-CasRx-*Vegfa* into one eye, and AAV-CasRx with no gRNA into the other eye as control (Fig. 1m and Supplementary Fig. 4c). To evaluate the potential toxicity of AAV-CasRx-*Vegfa*-mediated gene knockdown, we performed electroretinography (ERG) recording in mice at one and two months after the subretinal injection. Our results showed that there is no significant change in the responses in mice injected with AAV-CasRx-*Vegfa* compared to that in mice injected with PBS (Supplementary Fig. 5a,b). In addition, we examined the expression level of opsin in the retina at around 1 month after AAV injection. We found that injection of AAV-CasRx-*Vegfa* did not affect the opsin-positive areas (Supplementary Fig. 5c). Together, these results suggest that AAV-CasRx-mediated *Vegfa* knockdown is a safe way to treat AMD.

In summary, our results demonstrate that AAV-mediated delivery of CasRx can potently knockdown *Vegfa* mRNA and suppress pathogenic CNV development in a mouse model of AMD, supporting the notion that RNA-targeting CRISPR system could be useful for therapeutic purposes. The small size of CasRx is suitable for packaging with multiple gRNAs in a single AAV vector for *in vivo* delivery. Notably, AAV-delivered CasRx has the potential for sustained corrective effects on protein expression for up to 2 years with a single injection^14^. The risks associated with mRNA editing could be lower than that of DNA editing, because of the existence of large number of transcripts, many of which may maintain normal functions. Thus CasRx knockdown approach could complement existing therapeutic strategies such as monoclonal antibodies, antisense oligonucleotides and DNA nuclease editing. Intriguingly, a recent study demonstrated that Cas13 showed potent activity against RNA viruses^15^. In the future, it is promising to examine whether CasRx could be used to inhibit the reproduction of recently emerged deadly RNA viruses such as 2019-nCoV, Ebola, MERS and Zika.

## Acknowledgements

We thank Drs. Mu-ming Poo for helpful discussions and insightful comments on this manuscript. We thank Wenqin Ying, Qifang Wang, Yiwen Zhang, Yanli Lu for technical assistance and valuable discussion. This work was supported by R&D Program of China (2018YFC2000100 and 2017YFC1001302), CAS Strategic Priority Research Program (XDB32060000), National Natural Science Foundation of China (31871502, 31522037), Shanghai Municipal Science and Technology Major Project (2018SHZDZX05), Shanghai City Committee of science and technology project (18411953700, 18JC1410100).

## Author contributions

CZ designed and performed experiments. XH, CT, WL and QB performed CNV experiments. SW perfomed ERG. YZ analyzed the RNA-seq data. LS prepared AAVs. XS, HZ and HY designed experiments and supervised the project. HZ and HY wrote the paper.

## Methods

### Ethical compliance

The use and care of animals complied with the guideline of the Biomedical Research Ethics Committee of Institute of Neuroscience, Chinese Academy of Sciences.

### Vector and gRNA sequences

The vector information are provided in Supplementary Sequences. gRNA1: 5’-gtgctgtaggaagctcatctctcctatgtg-3’; gRNA2: 5‘-ggtactcctggaagatgtccaccagggtct-3’.

### Transient transfection, qPCR and RNA-seq

Plasmids transient transfection was performed as previously described^1^. 293T and N2a cells were cultured in dulbecco’s modified eagle medium (DMEM) containing 10% fetal bovine serum (FBS) and penicillin/streptomycin, and maintained at 37 °C with 5% CO_2_. Cells were seeded in 6-well plates and transfected with 4 μg/well vectors expressing CasRx-GFP and gRNAs-mCherry (CasRx:gRNA-1:gRNA-2 = 2:1:1, see Supplementary sequences) using Lipofectamine 3000 reagent (Thermo Fisher Scientific). Control group was only transfected with 2 μg/well vectors containing CasRx-GFP. GFP^+^mCherry^+^ cells (GFP^+^ cells for control group) were isolated using flow cytometry 3 days after transfection. Total RNA was first purified using Trizol (Ambion) and then transcribed into complementary DNA (HiScript Q RT SuperMix for qPCR, Vazyme, Biotech). qPCR reactions were tracked by SYBR green probe (AceQ qPCR SYBR Green Master Mix, Vazyme, Biotech). For Supplementary Fig. 3, DNA was extracted from the RPE complex and used for qPCR. VEGFA qPCR primers are: Forward, 5‘-GGTGGACATCTTCCAGGAGT-3’; Reverse, 5’-TGATCTGCATGGTAGATGTTG-3’.

CasRx qPCR primers are: Forward, 5‘-CCCTGGTGTCCGGCTCTAA-3’; Reverse, 5’-GGACTCGCCGAAGTACCTCT-3’. For RNA-seq, around 250000 GFP-positive cells were collected and lysed. Total RNAs were extracted and then converted to cDNA, which was used for RNA-seq. The libraries were sequenced using Illumina Xten platform. Low-quality reads were filtered with SolexaQA (V3.1.7.1) and aligned to mm10 reference genome with Hisat2 (V2.0.4). Read counts and differentially expressed genes were calculated using htseq-count (v0.11.2) and DEseq2 (1.24.0), respectively. Genes were treated as differentially expressed genes when the fold-change > 2 and FDR < 0.05.

### AAV production and intravitreal injection

AAV-CasRx-*Vegfa* and AAV-CasRx (AAV-PHP.eb capsid^2^) was packaged by transfection of HEK293T cells using Polyethylenimine (PEI) (50 μg/ml). Viruses were harvested, purified and concentrated 3–7 days after transfection. Mice (C57BL/6) aged 6-8 weeks were anesthetized for intravitreal injection. Intravitreal injection of PBS, AAV-CasRx or AAV-CasRx-*Vegfa* (7.5*10^9 viral genomes in 1 μl) was intravitreally injected using a Hamilton syringe with a 34G needle under an Olympus microscope (Olympus, Tokyo, Japan). Mice with retinal hemorrhage were excluded.

### Laser-induced CNV model, CNV staining and ERG

At 2-3 weeks after AAV injection, mice were used for laser burn. CNV models were induced as previously described^3^. In brief, mice were anesthetized and pupils were dilated with dilating eye drops to enlarge the pupil size. Laser photocoagulation was performed using NOVUS Spectra (LUMENIS). The laser parameters used in this study were: 532 nm wave length, 70 ms exposure time, 240 mW power (otherwise stated) and 50 μm spot size. 4 laser burns (30 laser burns for ELISA) around the optic disc were induced. Mice with vitreous hemorrhage were excluded in the study. 3 days after laser induction, mice were perfused with saline and RPE complexes were dissociated for ELISA analysis (qPCR was performed 7 days after laser burn). CNV analysis was conducted 7 days later after laser burn. Mice were perfused with PFA and the eyes were then fixed with PFA for 2 hours. The retina was removed from the eyes, and only RPE/choroid/scleral complex was stained with isolectin-B4 (IB4, 10μg/ml, I21413, Life Technologies) overnight. RPE complexes were flat-mounted and visualized with microscope (VS120 Olympus). Only eyes with successful AAV-CasRx-*Vegfa* infection were included for quantification. After obtaining CNV images, DNA was extracted from the RPE complex, the copy number of CasRx was evaluated by qPCR. The area of CNV was quantified using ImageJ software by a blind observer. ERG was performed as previously described^4^.

### ELISA

RPE complex were collected for ELISA. To perform VEGFA ELISA, 30 laser burns were induced in each eye 3 weeks after AAV injection. Eyes were enucleated 3 days post-induction, the RPE complexes were dissociated from the retinas and lysed with RIPA lysis buffer. VEGFA protein levels were determined using Quantikine ELISA kit (MMV00, R&D SYSTEMS) according to the standard protocol.

### Statistical analysis

All values are shown as mean ± s.e.m. Statistical significance (*p* < 0.05) is determined by unpaired two-tailed Student’s *t* test. Randomization was used in all experiments and no statistical methods were used to pre-determine sample sizes but our sample sizes are similar to those reported in previous publications^5^.

## Supplementary Figures

**Supplementary Figure 1.**
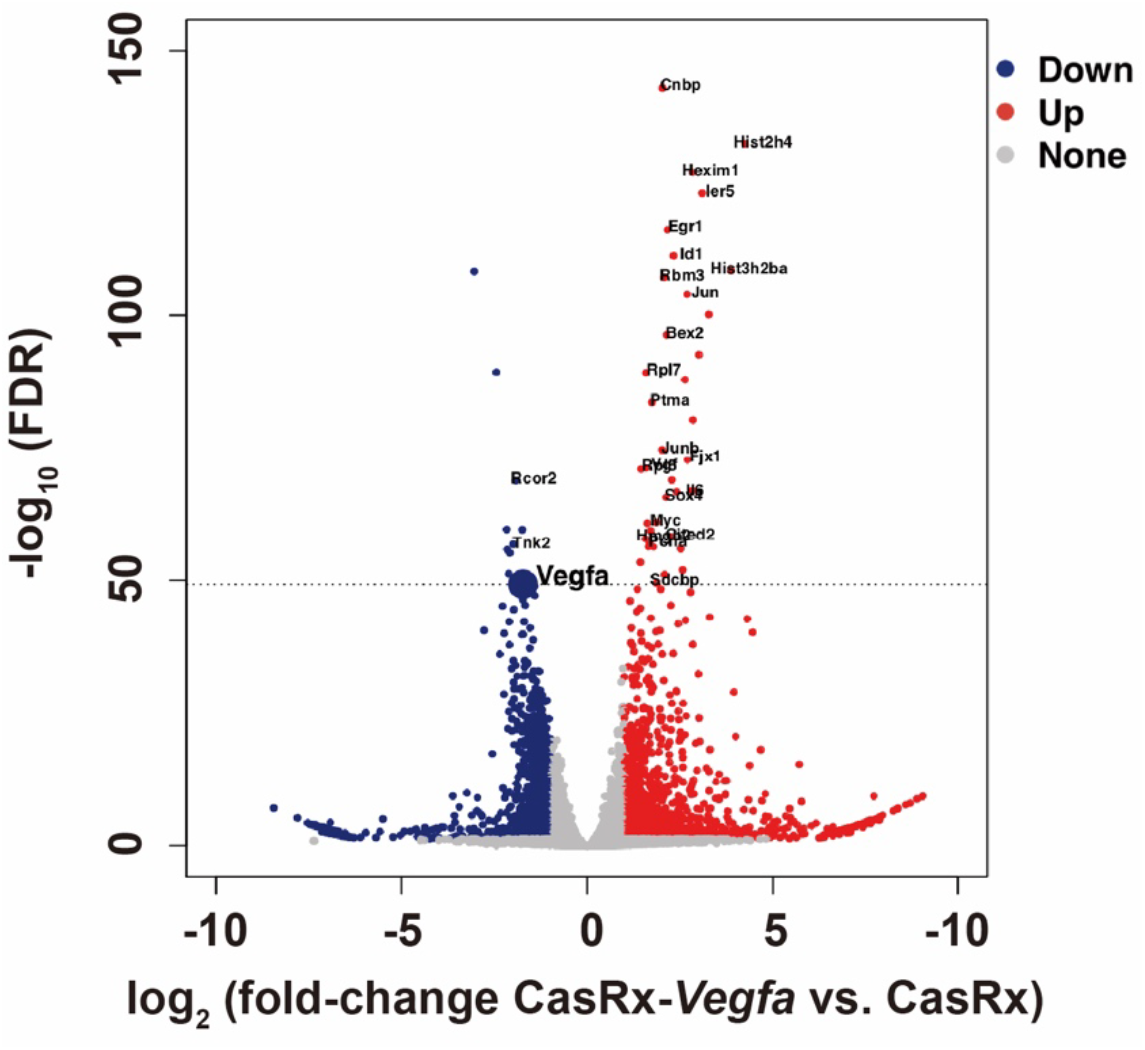
Targeting specificity of CasRx-*Vegfa*. Volcano plot showing the expression levels of all detected genes in RNA-seq libraries of CasRx-*Vegfa* compared to CasRx control. N2a cells, n = 3 independent replicates for both groups. Note that 24 out of 40 top-ranked genes with altered expression were related with *Vegfa* according to previous studies, and these genes are marked in the Figure. For details, see Supplementary Table 1.

**Supplementary Figure 2.**
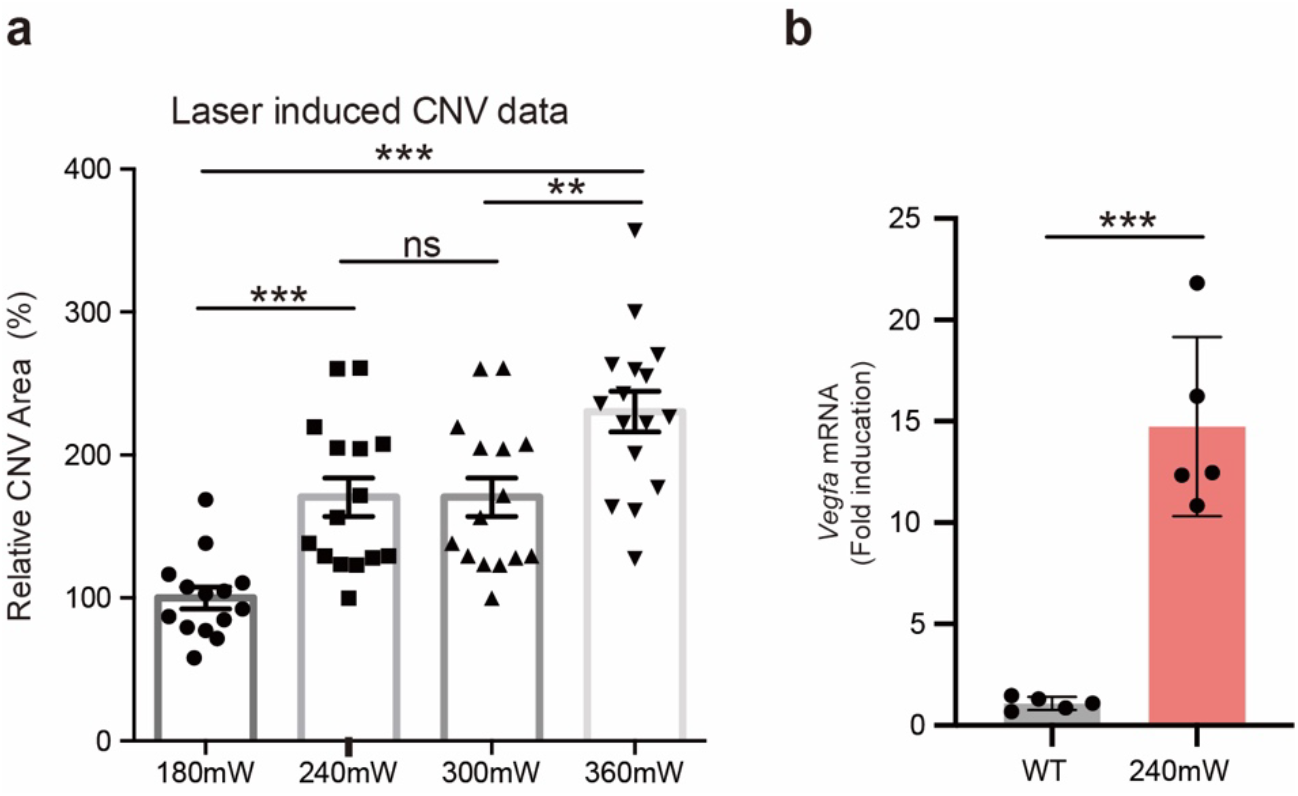
Induction of CNV with different levels of laser irradiation. (**a**) A data point represents a laser burn and in total 4 laser burns were induced in each eye. (**b**) Increase of *Vegfa* expression 7 days after laser burn (n =. 5 retinas per group). All values are presented as mean ± s.e.m.. \**p* < 0.05, \*\**p* < 0.01, \*\*\**p* < 0.001, unpaired t-test.

**Supplementary Figure 3.**
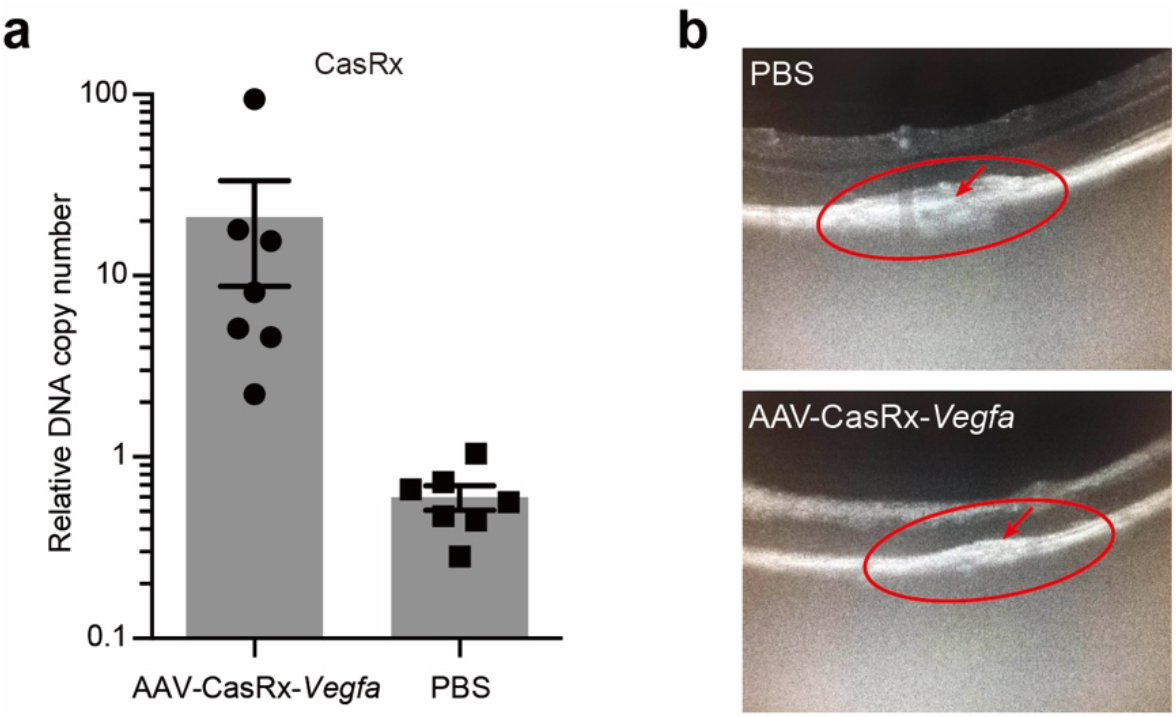
AAV expression and induction of CNV. **(a)** Successful infection of AAV-CasRx-*Vegfa* in the RPE complex after laser burn. **(b)** Images showing laser-induced CNV (red circle) in the anesthetized mice, 7 days after laser burn. All values are presented as mean ± s.e.m..

**Supplementary Figure 4.**
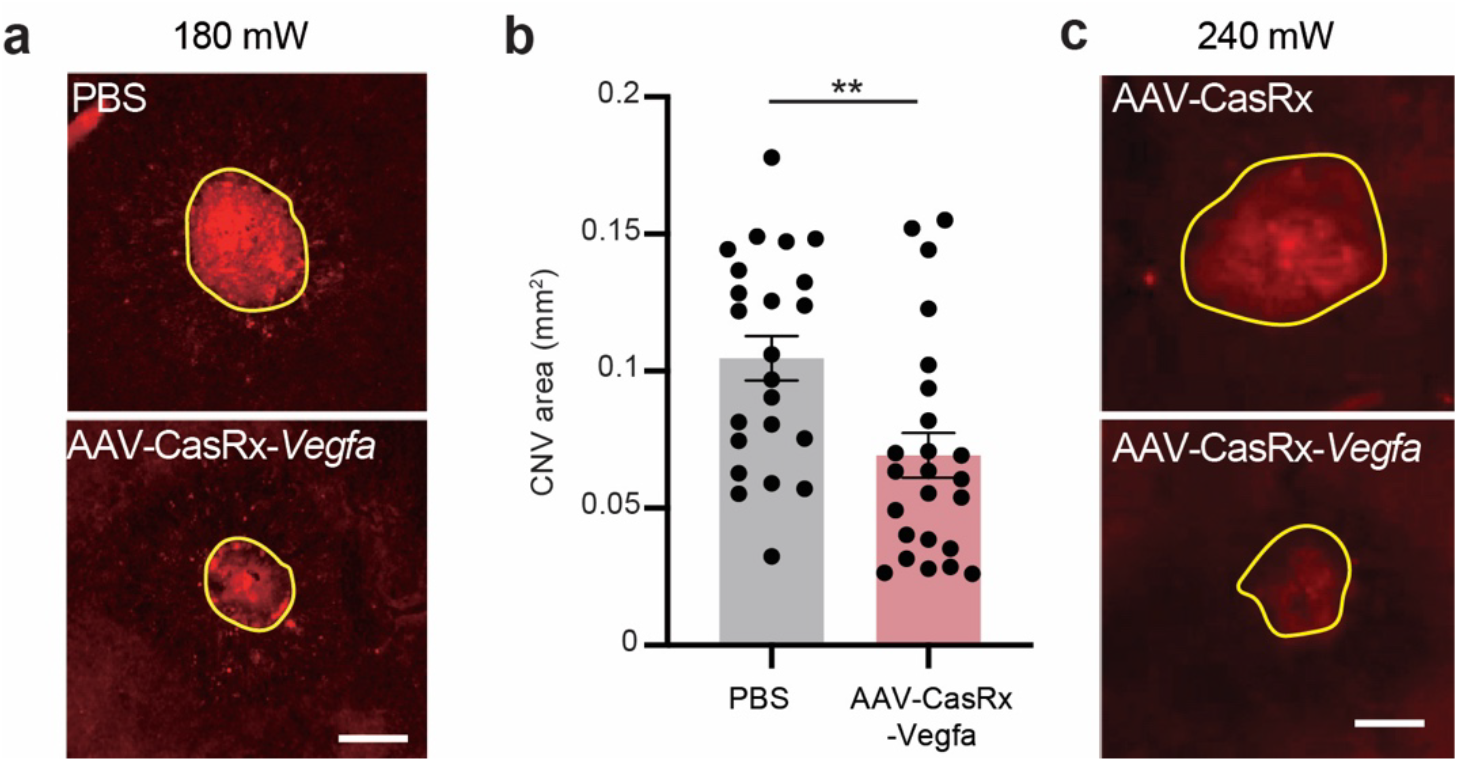
Reduction of the area of CNV at two different levels of laser irradiation. (**a**) Representative images showing CNV was induced in the retina injected with PBS or AAV-CasRx-*Vegfa* using a laser intensity of 180 mW. The area of CNV is indicated by the yellow line. Scale bar: 200 μm. (**b**) The CNV area. A data point represents a laser burn and in total 4 laser burns were induced in each eye. (180 mW, *n* = 6 mice, *p* = 0.004, *t* = 3.079). (**c**) Representative images showing CNV was induced in the retina injected with AAV-CasRx or AAV-CasRx-*Vegfa* using a laser intensity of 240 mW. The area of CNV is indicated by the yellow line. Scale bar: 200 μm. All values are presented as mean ± s.e.m.. \**p* < 0.05, \*\**p* < 0.01, \*\*\**p* < 0.001, unpaired t-test.

**Supplementary Figure 5.**
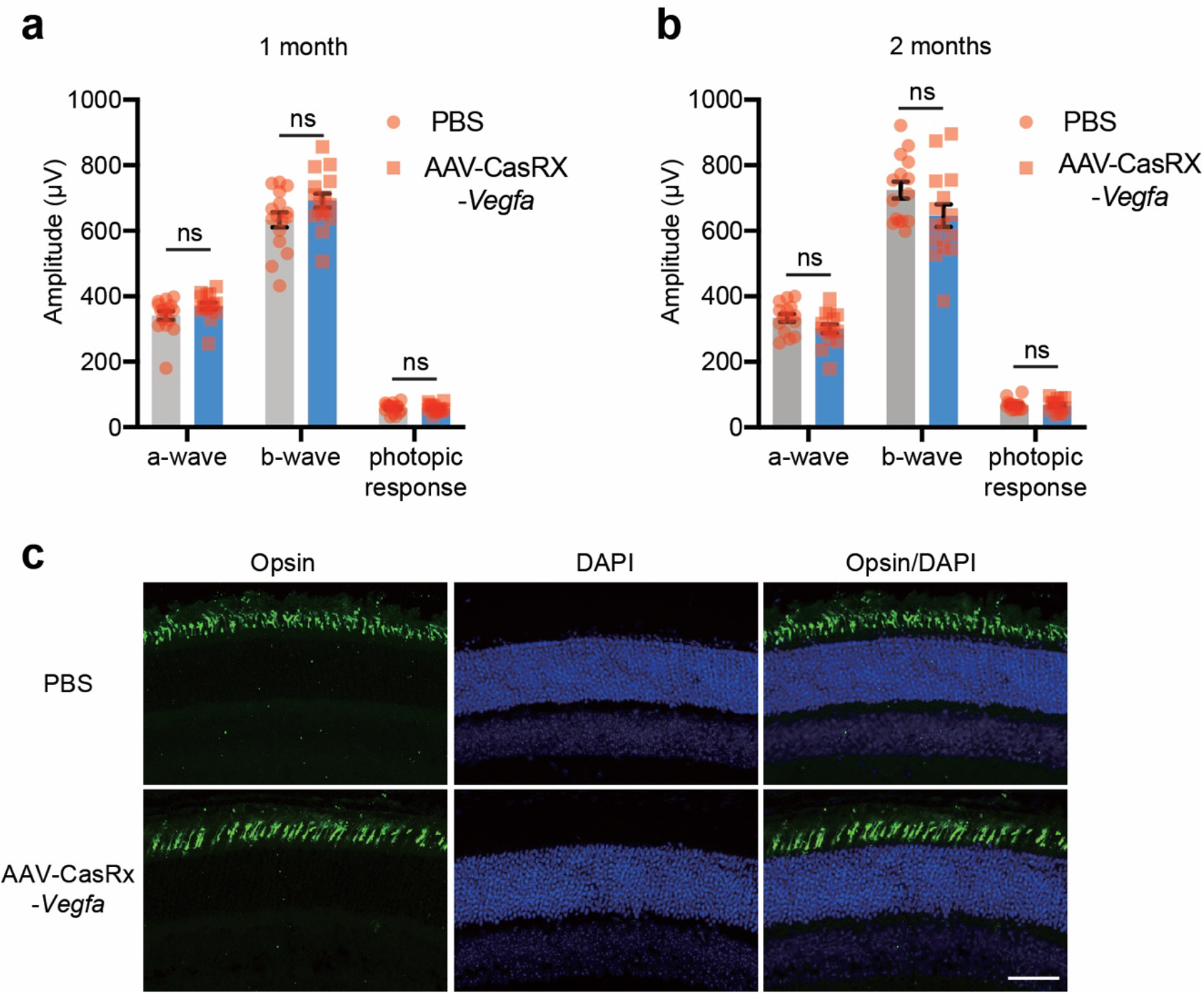
Injection of AAV-CasRx-*Vegfa* does not affect the retinal function. (**a,b**) ERG was performed to evaluate retinal function. Full-field ERG showed that there was no significant change in the scotopic a-wave, scotopic b-wave, or photopic response in mice injected with AAV-CasRx-*Vegfa* compared to control mice injected with PBS at 1 or 2 months after injection (1 month: n = 16 mice per group; 2 month: n = 15 mice per group). (**c**), Representative images showing that the size of opsin-positive area was not affected 1 month after injection (n = 3 mice). Scale bar: 50 μm. All values are presented as mean ± s.e.m.. \**p* < 0.05, \*\**p* < 0.01, \*\*\**p* < 0.001, unpaired t-test.

## Supplementary Tables

**Supplementary Table 1. List of all genes from RNA-seq.** Note that 24 out of 40 top-ranked genes with altered expression (based on p value, yellow) were reported related with *Vegfa*.

## Supplementary Sequences

### CasRX-P2A-GFP plasmid

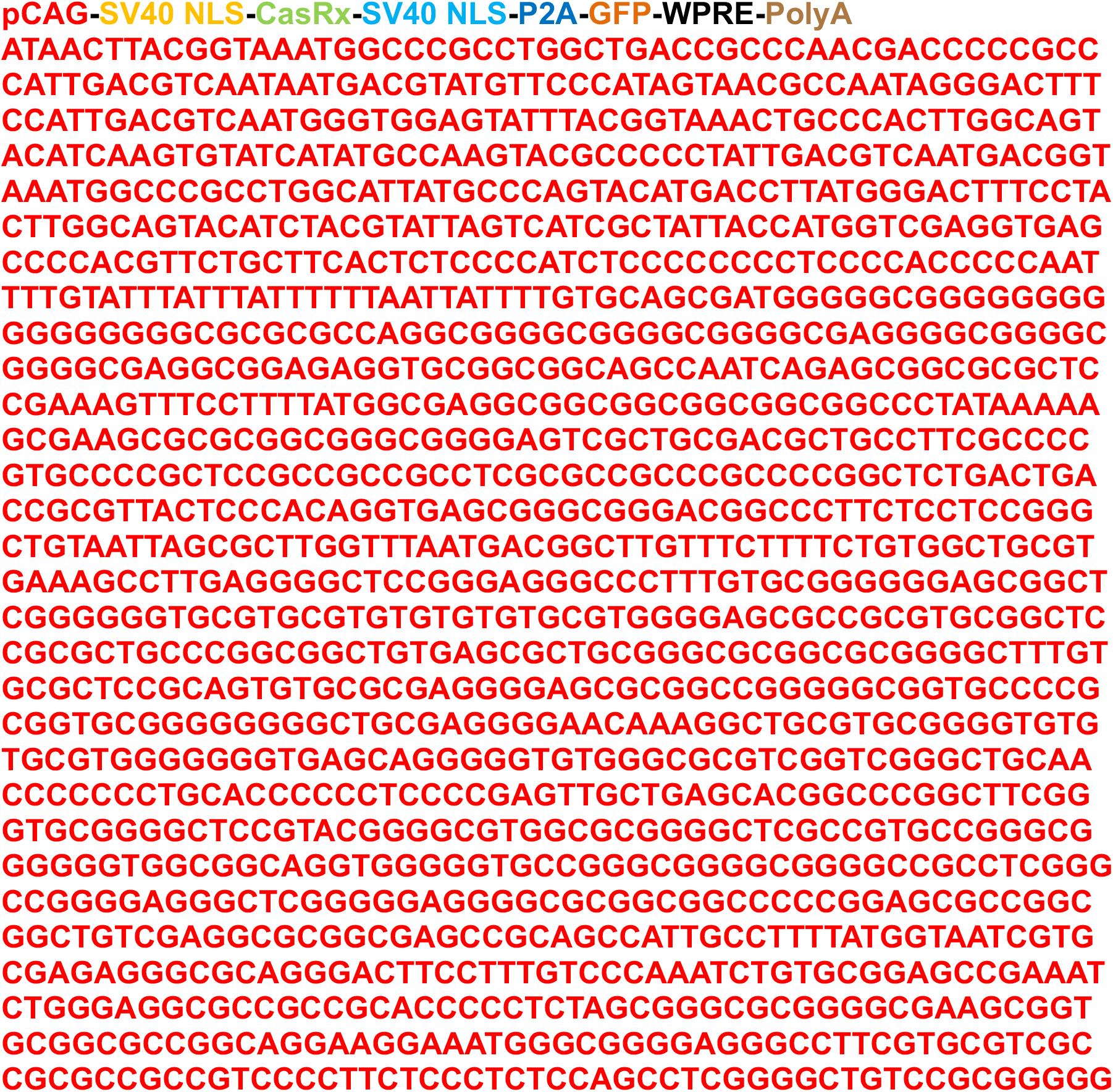
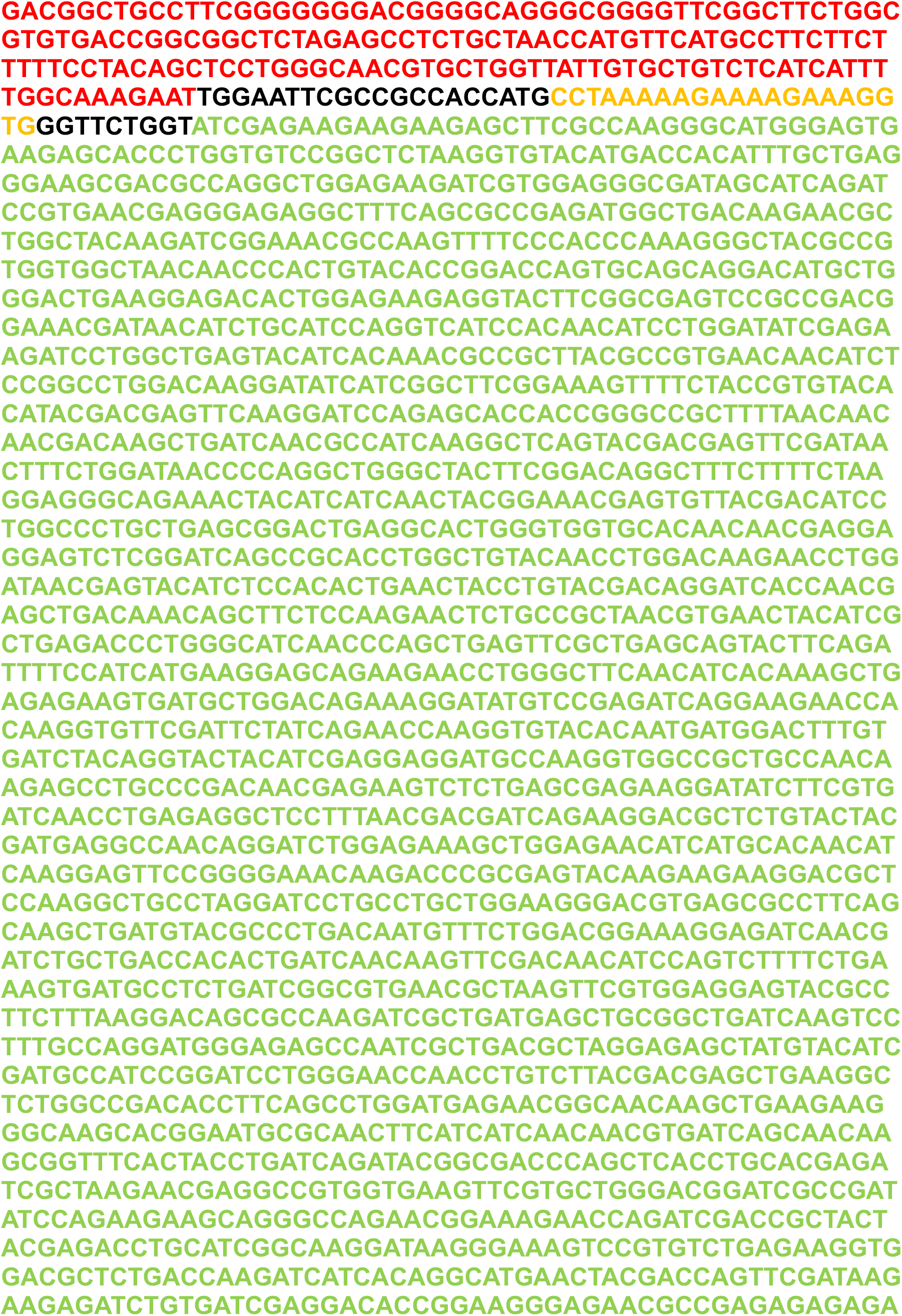
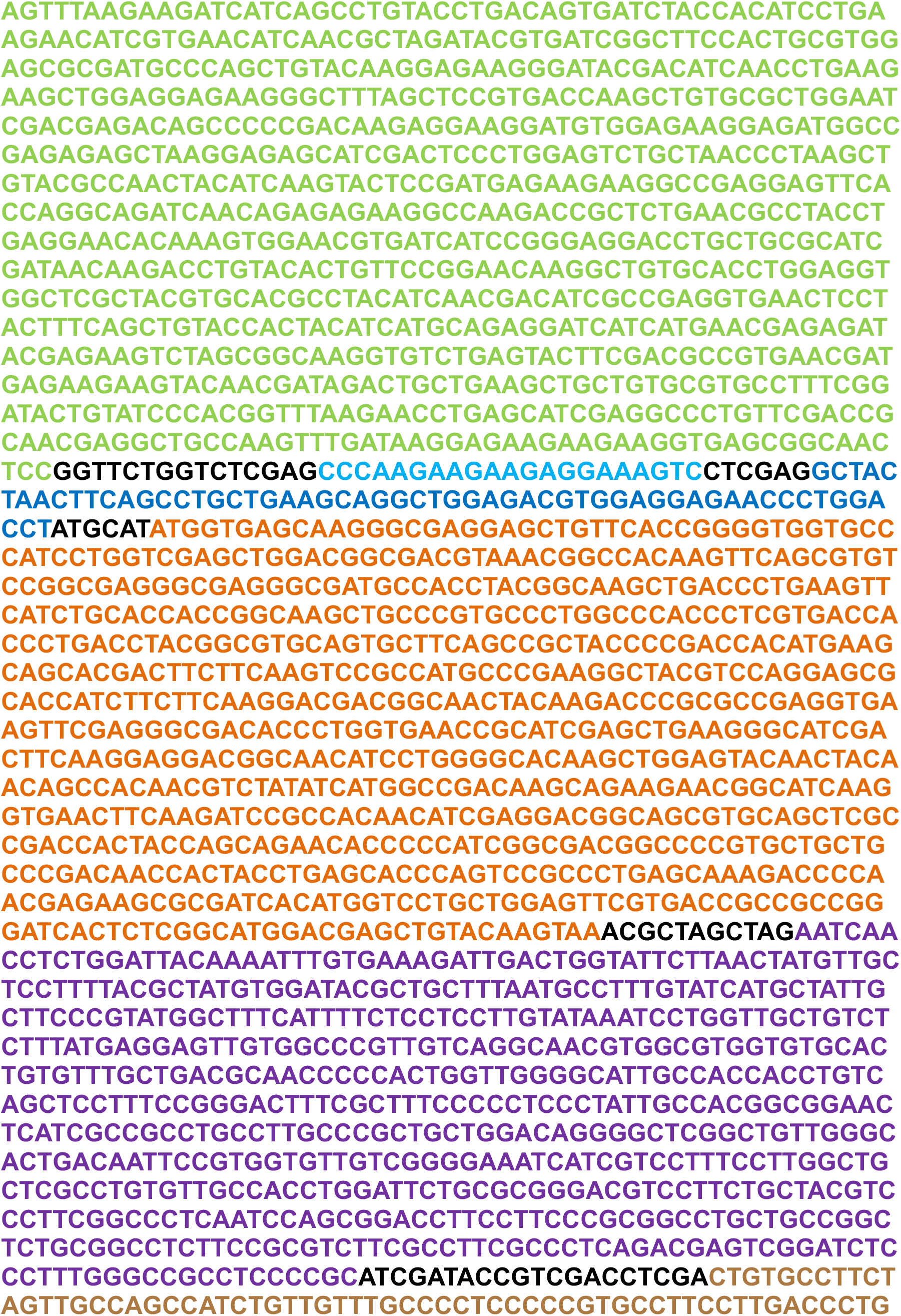
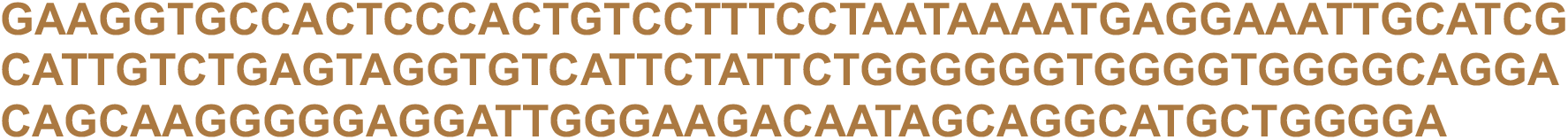

### Vegfa-gRNA-1 plasmid

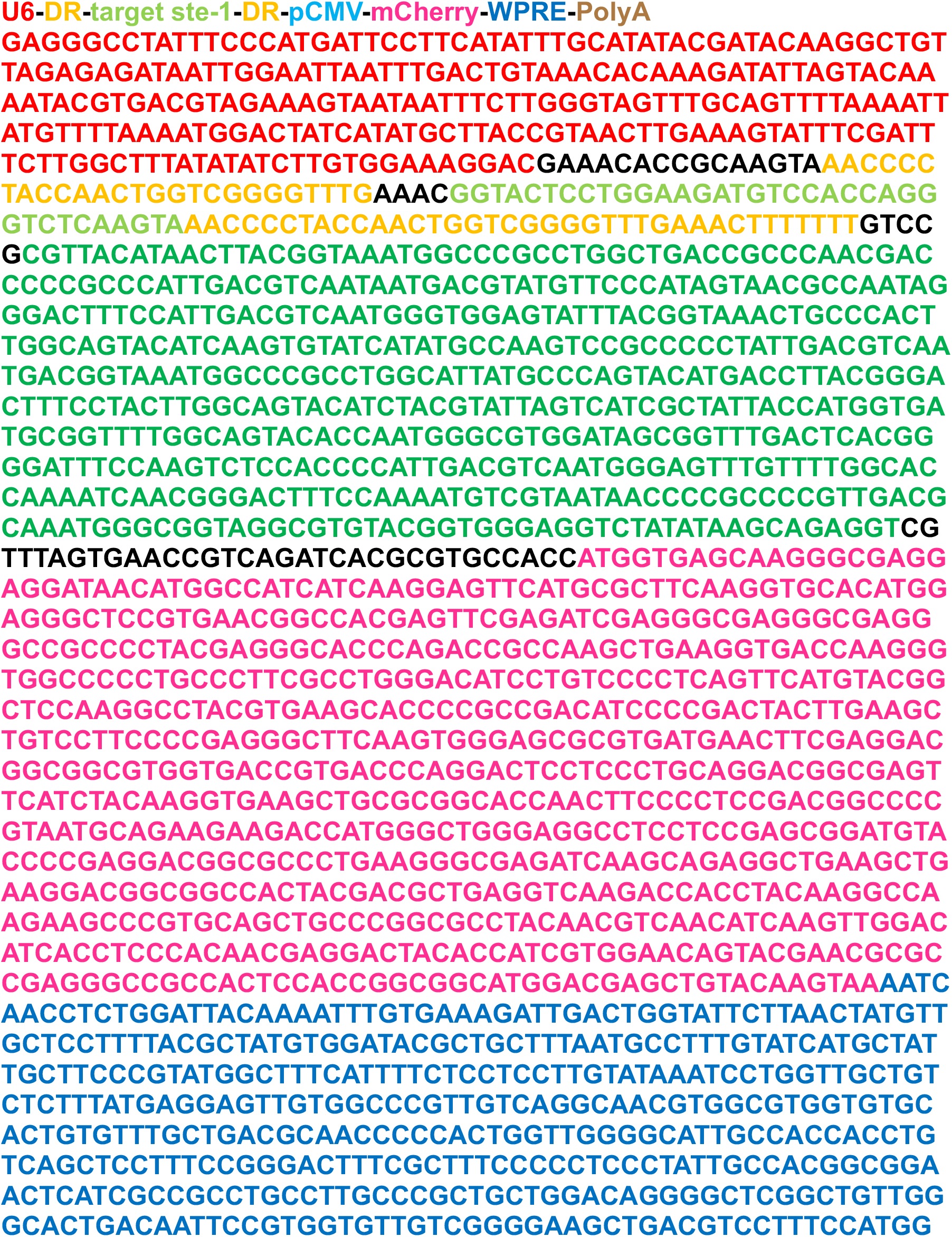
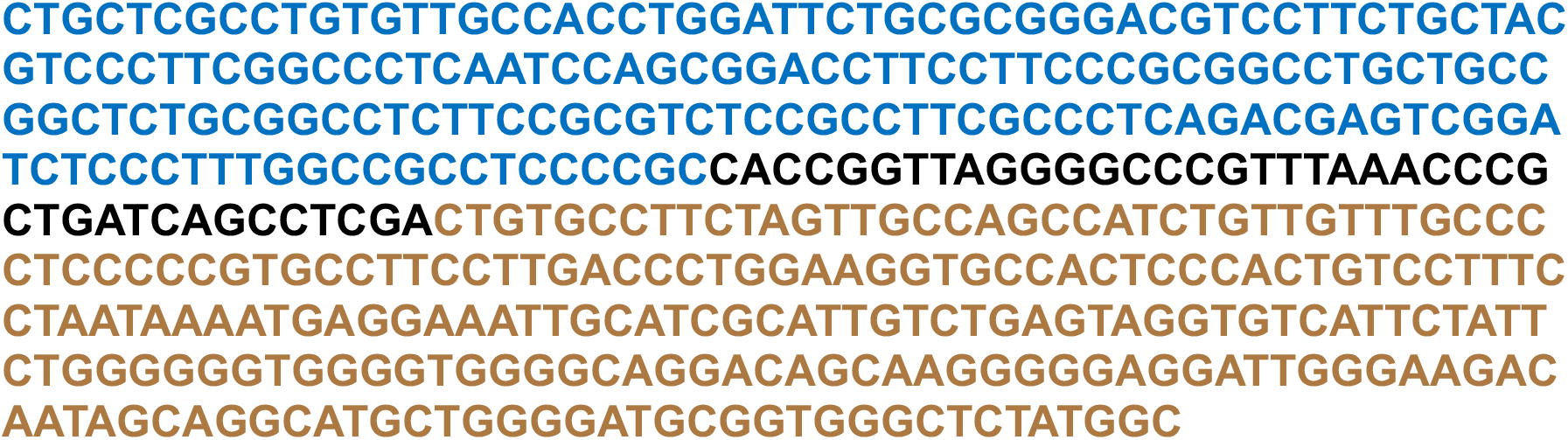

### Vegfa-gRNA-2 plasmid

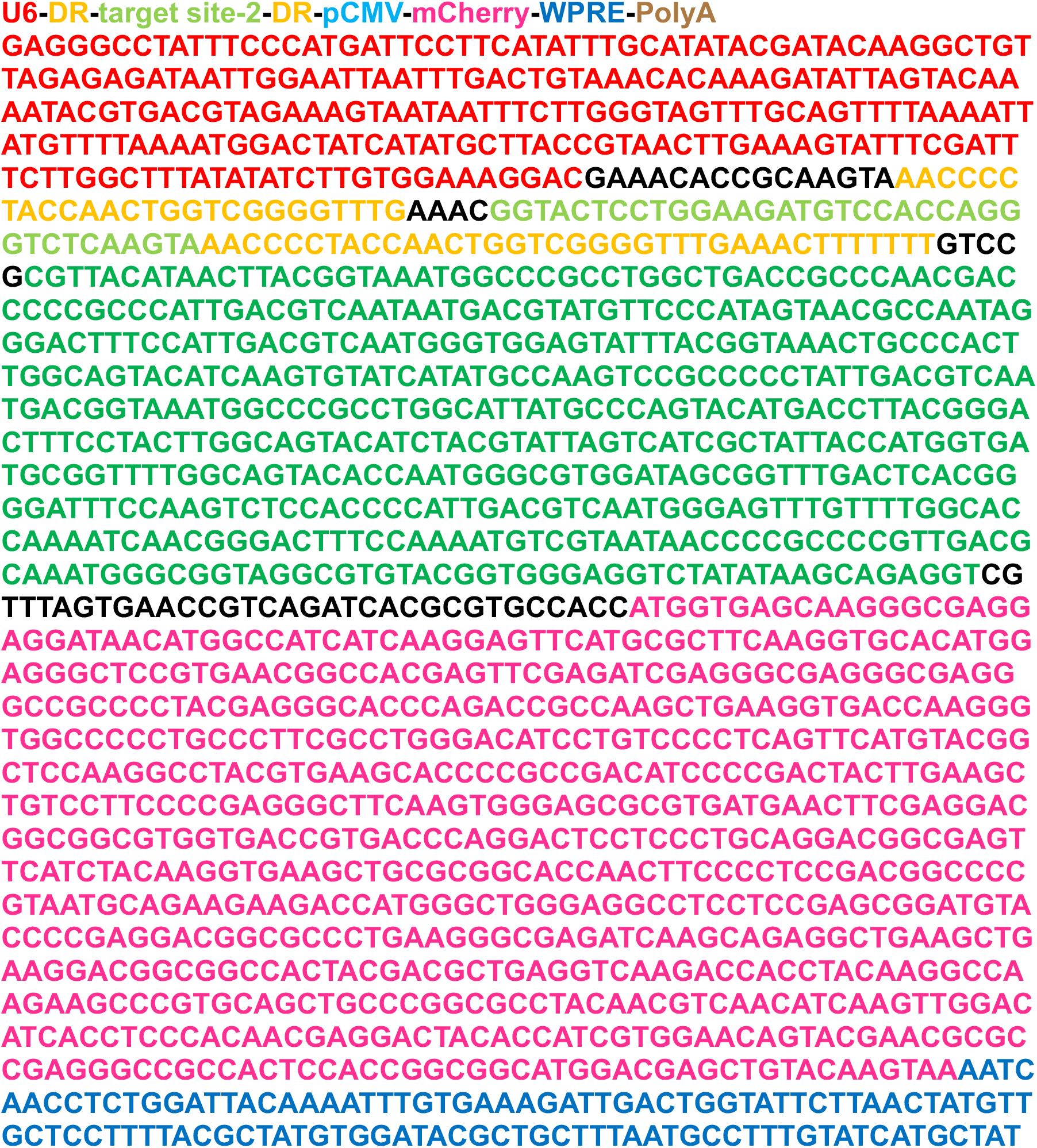
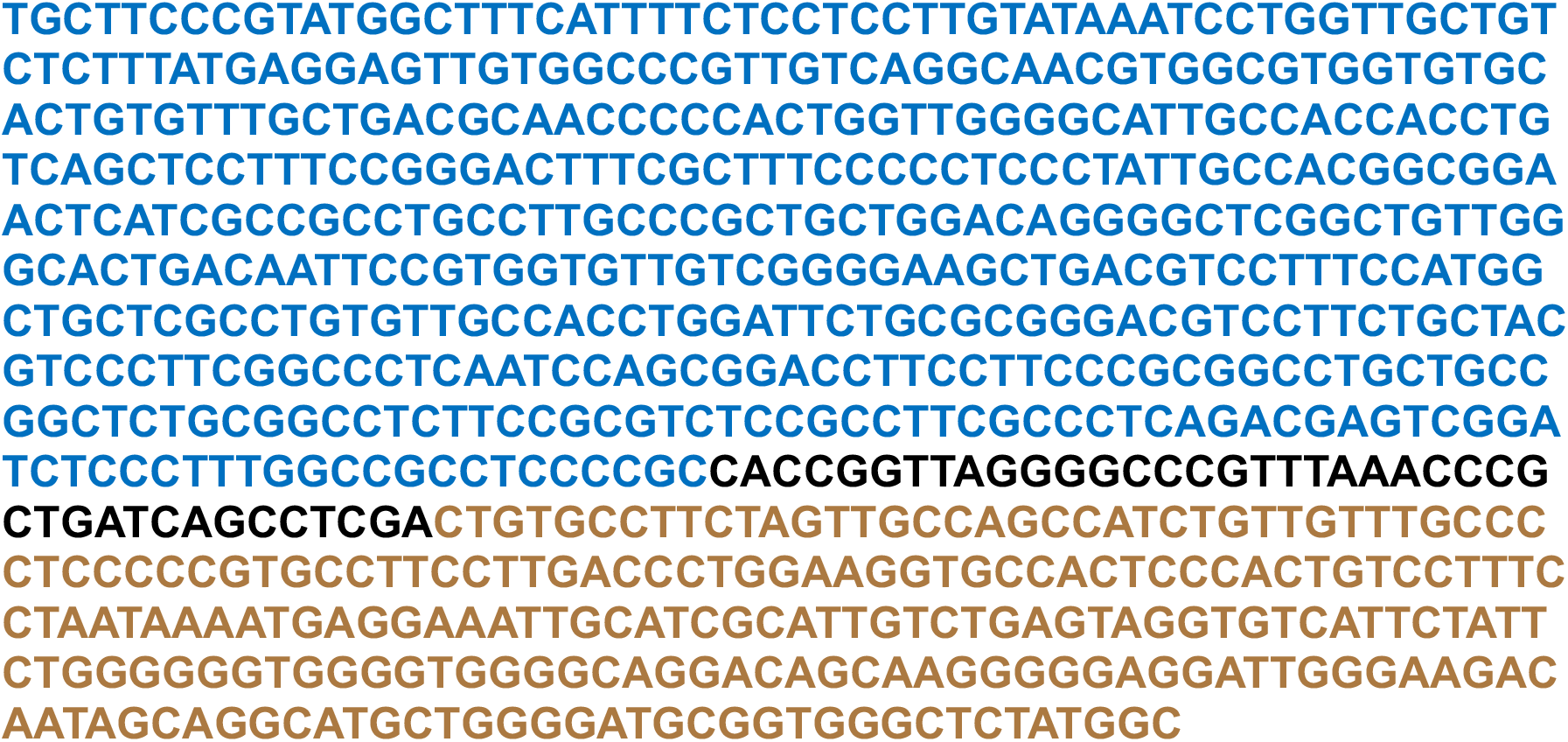

### AAV-CasRx-Vegfa plasmid

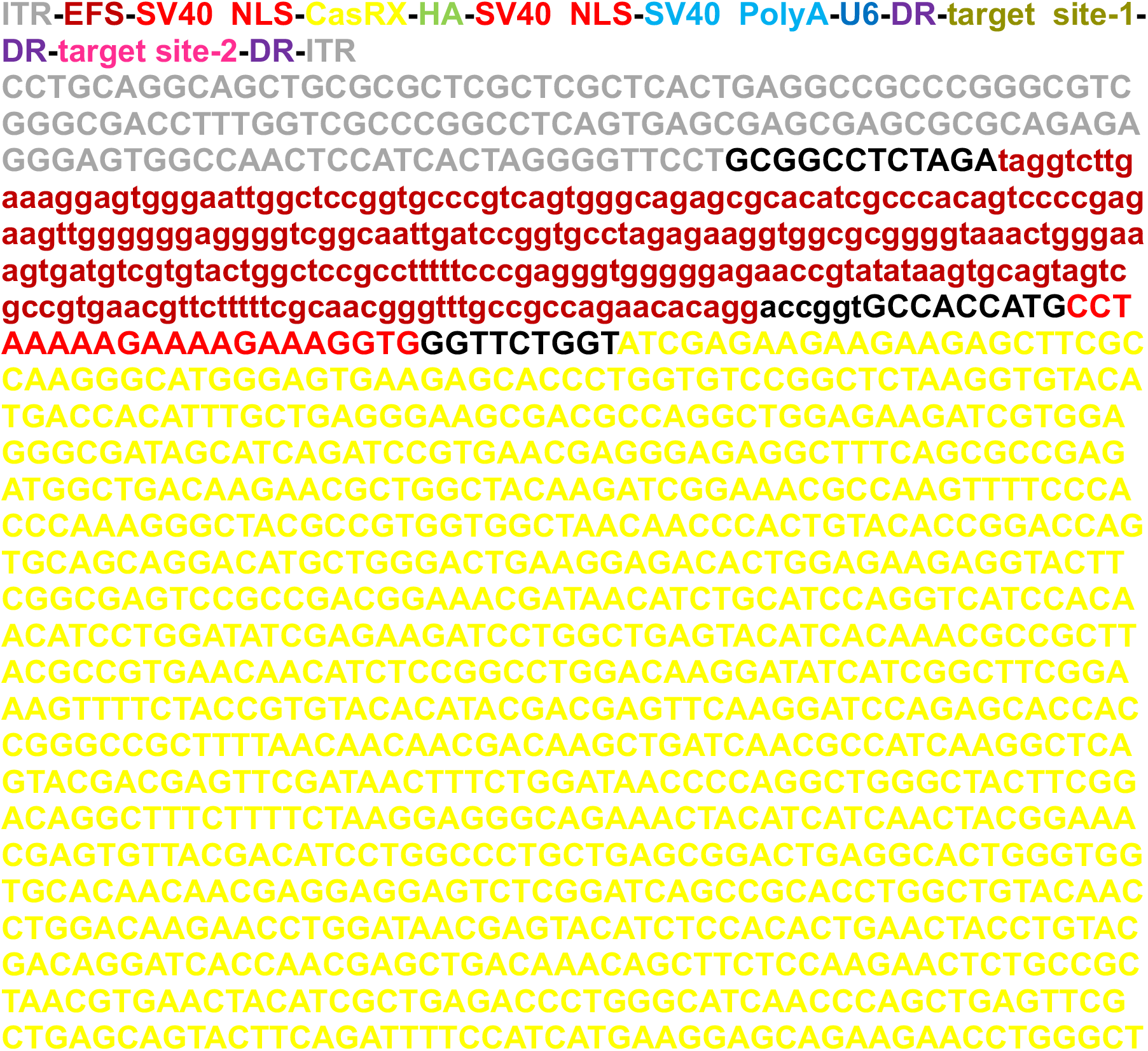
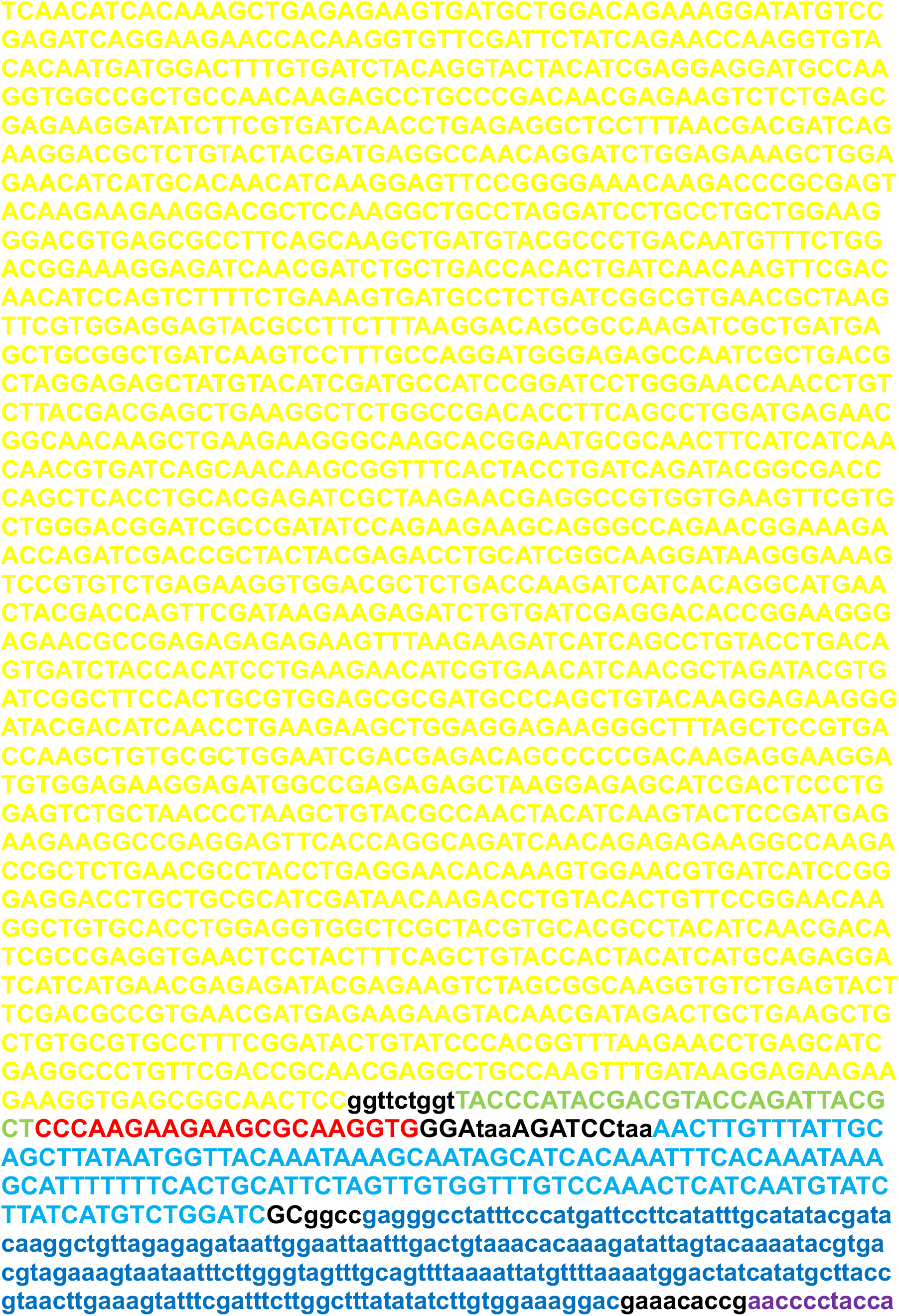
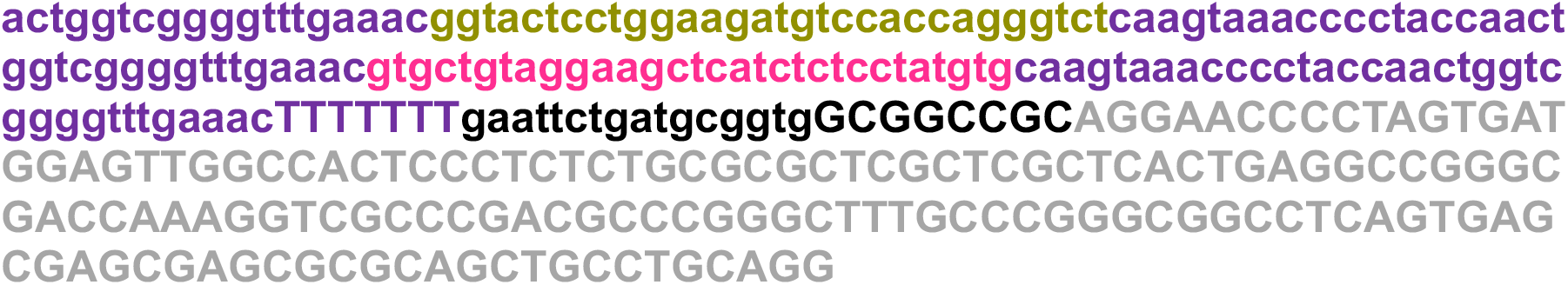

### AAV-CasRx plasmid

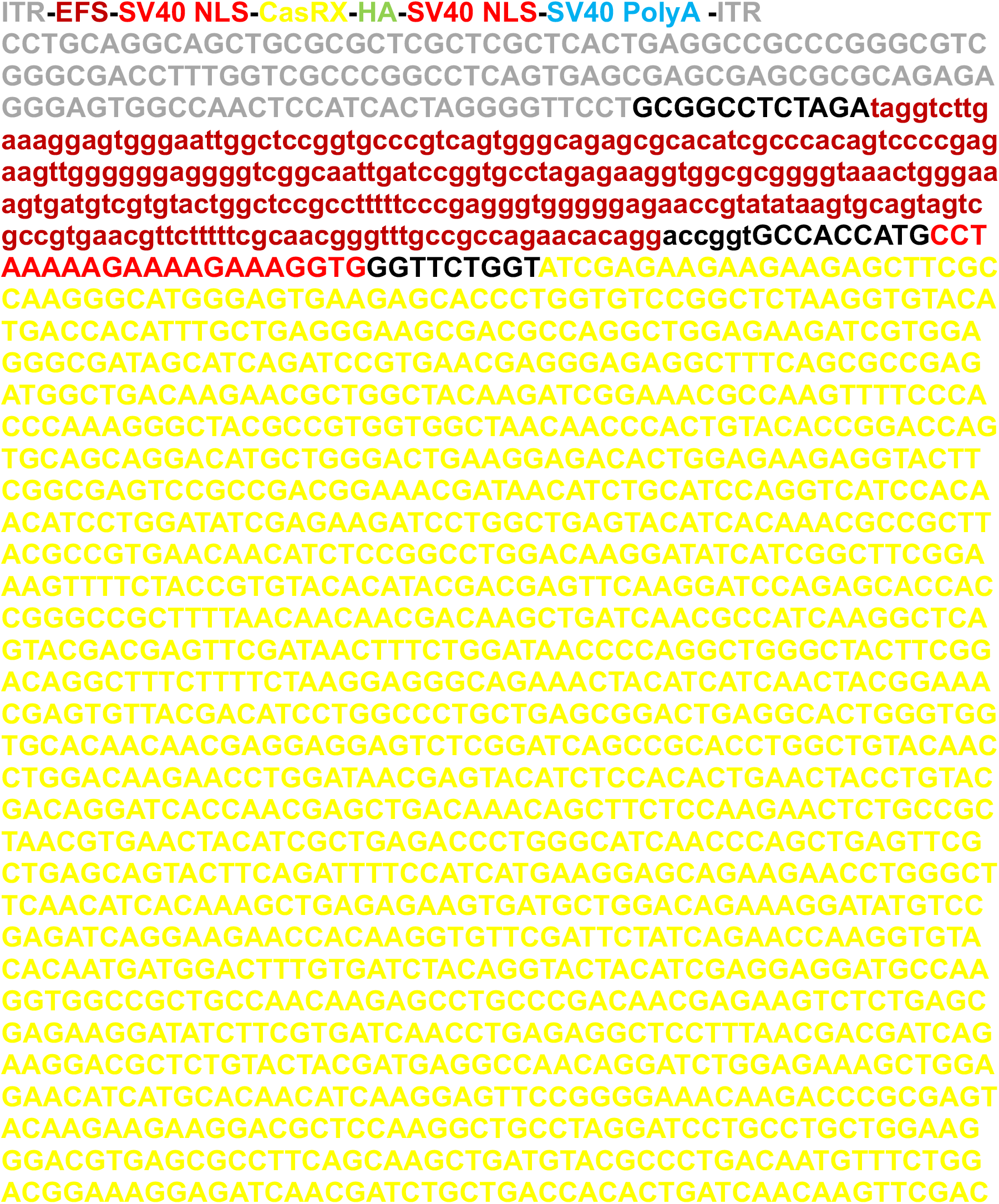
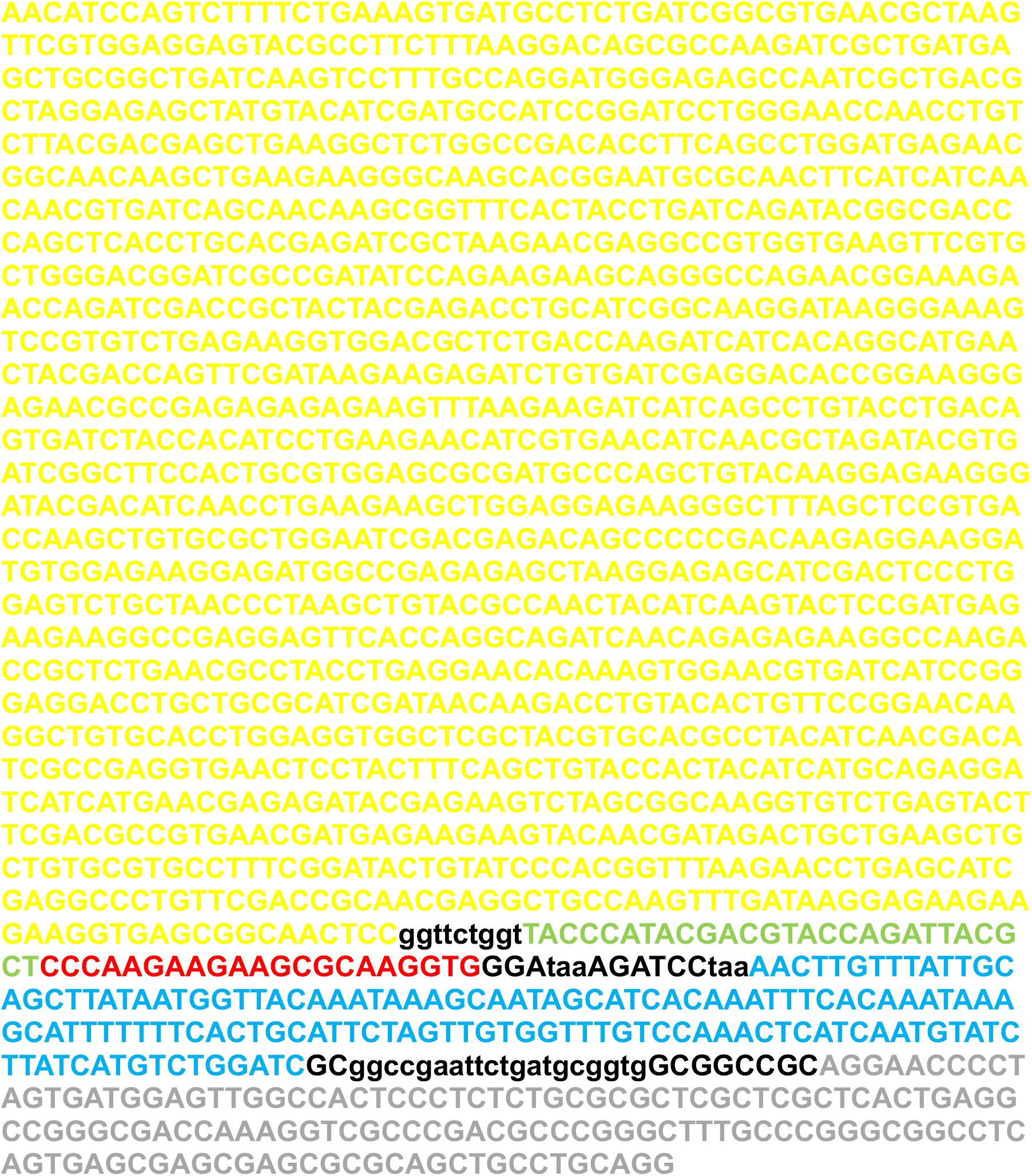

## References

1 Cheung, L. K. & Eaton, A. Age-related macular degeneration. Pharmacotherapy 33, 838–855, doi:10.1002/phar.1264 (2013).

2 Amoaku, W. M. et al. Defining response to anti-VEGF therapies in neovascular AMD. Eye 29, 721–731, doi:10.1038/eye.2015.48 (2015).

3 Ferrara, N. Vascular endothelial growth factor and age-related macular degeneration: from basic science to therapy. Nat Med 16, 1107–1111, doi:10.1038/nm1010-1107 (2010).

4 Kim, K. et al. Genome surgery using Cas9 ribonucleoproteins for the treatment of age-related macular degeneration. Genome Res 27, 419–426, doi:10.1101/gr.219089.116 (2017).

5 Koo, T. et al. CRISPR-LbCpf1 prevents choroidal neovascularization in a mouse model of age-related macular degeneration. Nat Commun 9, doi:Artn185510.1038/S41467-018-04175-Y (2018).

6 Kosicki, M., Tomberg, K. & Bradley, A. Repair of double-strand breaks induced by CRISPR-Cas9 leads to large deletions and complex rearrangements (vol 36, pg 765, 2018). Nature Biotechnology 36, 899–899, doi:Doi 10.1038/Nbt0918-899c (2018).

7 Shin, H. Y. et al. CRISPR/Cas9 targeting events cause complex deletions and insertions at 17 sites in the mouse genome. Nat Commun 8, doi:Artn 15464 10.1038/Ncomms15464 (2017).

8 Abudayyeh, O. O. et al. RNA targeting with CRISPR-Cas13. Nature 550, 280-+, doi:10.1038/nature24049 (2017).

9 Cox, D. B. T. et al. RNA editing with CRISPR-Cas13. Science 358, 1019–1027, doi:10.1126/science.aaq0180 (2017).

10 East-Seletsky, A. et al. Two distinct RNase activities of CRISPR-C2c2 enable guide-RNA processing and RNA detection. Nature 538, 270-+, doi:10.1038/nature19802 (2016).

11 Knott, G. J. & Doudna, J. A. CRISPR-Cas guides the future of genetic engineering. Science 361, 866–869, doi:10.1126/science.aat5011 (2018).

12 Konermann, S. et al. Transcriptome Engineering with RNA-Targeting Type VI-D CRISPR Effectors. Cell 173, 665-+, doi:10.1016/j.cell.2018.02.033 (2018).

13 Shmakov, S. et al. Discovery and Functional Characterization of Diverse Class 2 CRISPR-Cas Systems. Molecular Cell 60, 385–397, doi:10.1016/j.molcel.2015.10.008 (2015).

14 Mendell, J. R. et al. Single-Dose Gene-Replacement Therapy for Spinal Muscular Atrophy. New Engl J Med 377, 1713–1722, doi:10.1056/Nejmoa1706198 (2017).

15 Freije, C. A. et al. Programmable Inhibition and Detection of RNA Viruses Using Cas13. Mol Cell 76, 826–837 e811, doi:10.1016/j.molcel.2019.09.013 (2019).

## References

1 Zhou, H. et al. In vivo simultaneous transcriptional activation of multiple genes in the brain using CRISPR-dCas9-activator transgenic mice. Nat Neurosci 21, 440–446, doi:10.1038/s41593-017-0060-6 (2018).

2 Chan, K. Y. et al. Engineered AAVs for efficient noninvasive gene delivery to the central and peripheral nervous systems. Nat Neurosci 20, 1172–1179, doi:10.1038/nn.4593 (2017).

3 Gong, Y. et al. Optimization of an Image-Guided Laser-Induced Choroidal Neovascularization Model in Mice. PLoS One 10, e0132643, doi:10.1371/journal.pone.0132643 (2015).

4 Koo, T. et al. CRISPR-LbCpf1 prevents choroidal neovascularization in a mouse model of age-related macular degeneration. Nat Commun 9, 1855, doi:10.1038/s41467-018-04175-y (2018).

5 Chavez, A. et al. Comparison of Cas9 activators in multiple species. Nat Methods 13, 563–567, doi:10.1038/nmeth.3871 (2016).

